# Growth and immunolocalisation of the brown alga *Ectocarpus* in a microfluidic environment

**DOI:** 10.1101/2021.07.20.453111

**Authors:** Bénédicte Charrier, Samuel Boscq, Bradley J. Nelson, Nino F. Läubli

## Abstract

PDMS chips have proven to be suitable environments for the growth of several filamentous organisms. However, depending on the specimen, the pattern of growth and cell differentiation has been rarely investigated. We monitored the developmental pattern of the brown alga *Ectocarpus* inside a PDMS lab-on-chip. Two main methods of inoculation of the lab-on-chip were tested, i.e. by injection of spores or by insertion of sporophyte filaments into the chamber. Growth rate, growth trajectory, cell differentiation, and branching were the main development steps that were monitored for 20 days inside 25 μm or 40 μm parallel channels under standard light and temperature conditions. They were shown to be similar to those observed in non-constrained *in-vitro* conditions. Labelling of *Ectocarpus* cell wall polysaccharides – both with calcofluor for cellulose, and by immunolocalisation for alginates with monoclonal antibodies–showed expected patterns when compared to open space growth using either epifluorescence or confocal microscopy. Overall this article describes the experimental conditions for observing and studying the basic unaltered processes of brown algal growth using microfluidic technology, which provides the basis for future biochemical and biological research.

## Introduction

Because of their highly polarised shape, filamentous organisms raise many questions, mainly concerning their growth. Is growth restricted to the apical cell, which corresponds to a case of localised growth, or is it shared by all the cells composing the filament as in cases of diffuse growth? How do intercellular transport and communication take place in such a polarised and uniaxial living material? What are the cues for cell differentiation that are distributed along such a linear tissue? How is the repetition of all these functions controlled when architectural complexity is increased through branching?

Several pieces of work have used microfluidic technologies to address these questions in a range of organisms spanning the evolutionary tree, including mammalian and plant cells (Peyrin et al., 2011; Tong et al., 2015; Siddique et al., 2013; Burri et al., 2018; Shamsudhin et al., 2017). Reasons for using microfluidics are numerous and reflect investigations requiring spatial constraints of the culture material, for example limiting the height and width of the available environment in the chip path to simplify long-term monitoring and improve resolution of microscopy-based observations (Kozgunova and Goshima, 2019) (Shamsudhin et al., 2016) (Zhou et al., 2021). It can also allow to separate filaments growing too densely otherwise (Bascom et al., 2016) or force them to grow perpendicular to a measuring device positioned at the exit of the chip (e.g. force sensor; Burri et al., 2018). Constrained paths can also display the existence of directional memory of growing filaments (Held et al., 2011). In addition to their control over growth directions, microfluidic chips can provide automated flow of chemical compounds (Agudelo et al., 2013) and apply electrical pulses (Agudelo et al., 2016), both perceived as cues for the activitation or repression of downstream signalling pathways.

However, the potential of such devices must first be evaluated for each biological model to avoid misinterpretations of subsequent results due to unidentified effects induced by the constraining environment. In the protonemata of the moss *Physcomitrella*, a study first showed that growth rate, cell differentiation, protoplast regeneration, and responses to drugs were all observed as under standard growth conditions (Bascom et al., 2016), thereby validating the use of polydimethylsiloxane (PDMS) chips. However, the response to these conditions depends, to some extent, on the geometric parameters of the chip. The available height between the glass slide and the PDMS layer was shown to have an impact on certain cellular biological processes. When moss protonemata filaments were constrained (lab-on-chip height of 4.5 μm *vs*. 21 μm for protonemata diameter), microtubule velocity was reduced while the viability and growth of the filaments were not impaired (Kozgunova and Goshima, 2019). In the fungus *Neurospora crassa*, growth in maze and grid PDMS meshworks was shown to significantly impair both growth rate and branching pattern (Held et al., 2011), but this was attributed to the chip pathway design rather than the PDMS polymer that is known to be chemically compatible and non-toxic.

Here, we evaluated the ability of *Ectocarpus sp*. to grow within PDMS-based microfluidic devices and assessed the suitability of our approach for immunocytochemistry. *Ectocarpus sp*. is a filamentous brown alga. Brown algae (Phaeophyceae) are a class of photo-autotrophic organisms that evolved independently of animals and plants. The ancestor of the kingdom they belong to, the Stramenopiles, diverged 1.6 billion years ago from the eukaryotic ancestor shared by the animal and plant lineages, Opisthokonta and Archaeplastida respectively (Baldauf, 2003). Since the knowledge of the first genomic sequence of brown algae (Cock et al., 2010), metabolic, cellular and developmental studies have confirmed their peculiar phylogenetic position, as they combine animal and plants characteristics (Bogaert et al., 2019; Bothwell et al., 2008; Cock et al., 2010; Michel et al., 2010; Popper et al., 2011, Rabille et al., 2019a). While *Ectocarpus* was shown to grow on PDMS surfaces immersed in seawater (Evariste et al., 2012), we studied how its filaments cope with the restricted and constrained spatial environment provided by a microfluidic chip. We focused on the prostrate filament of the sporophyte because it grows immediately after mitospore or zygote germination. More importantly, this filament grows by tip growth, a mechanism that is shared by many organisms belonging to other phyla of the eukaryotic evolution tree, thereby allowing macroevolutive scale comparisons. In addition to the oomycetes belonging to the Stramenopiles like brown algae do, tip growing cells include plant pollen tube and plant root hairs, moss filaments (protonemata) and algal rhizoid of the Archaeplastida phylum, and fungal hyphae and neurons in the Opisthokonta (Fig 1). Besides its different evolutionary history, *Ectocarpus* differs from other tip-growing organisms in many ways. Its sporophyte filaments are the slowest tip-growing organisms reported to date with a growth speed of 2.5 μm.h^−1^ (Fig 1), i.e. 700 times slower than the pollen tube (Rabillé et al., 2019a). In addition, we showed that the mechanisms of tip growth selected by this alga rely on a control of the cell wall thickness at the very tip of the apical cell, while most of the other tip growing cells, including the pollen tube, control growth by the variation of the cell wall stiffness at this position (Rabillé et al., 2019a). Further studies of tip growth mechanisms in *Ectocarpus*, namely those controlling the cell wall thickness during apical growth, requires the development of technologies allowing to monitore simultaneously several filaments growing in similar conditions over a long period of time.

**Figure 1:**
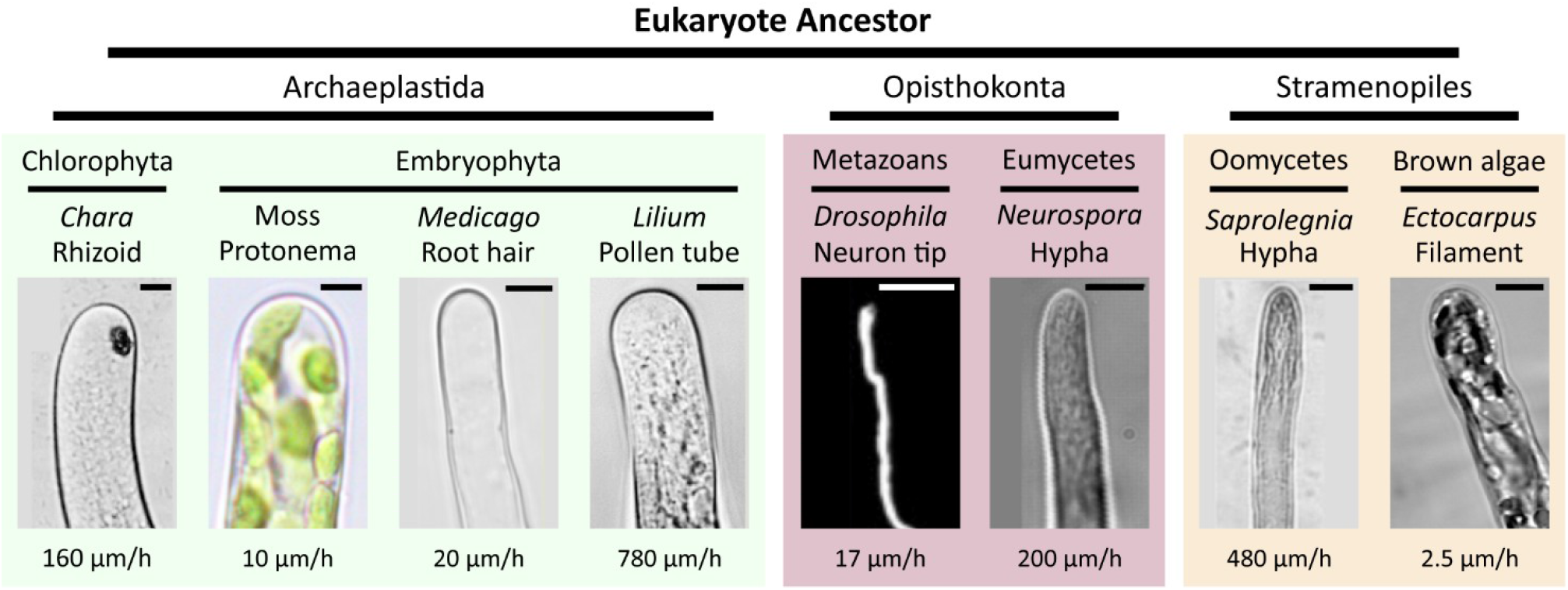
Diversity of tip growth in the Eukaryotic tree. Phylogenetic position of eukaryotic taxa with tip-growing organisms. Cell shapes and growth rates are shown. (A, B, C, D) Archaeplastida group: green algal filament, moss protonema, root hair; pollen tube. Stramenopiles: coenocytic oomycetes and the filamentous brown alga *Ectocarpus*; Opisthokonta group: neurons of metazoans and fungal hyphae. Scale bar = 5 μm (A, B, C, E, H, I, J), 10 μm (G), 20 μm (D, F). Modified from Rabillé et al. (2019a).

In this technical paper we designed two chips consisting of parallel channels, the purpose of which is to constrain the filaments to grow in straight directions parallel to each other, rather than in all directions outward as in open space. The healthiness and growth performance of *Ectocarpus* filaments in these conditions were monitored for several days and the growth rate was quantitatively compared to open space growth. In addition, the ability to label cytological markers, either by immunochemistry or directly with vital dyes, was tested. We showed that all the developmental steps tested during the growth of *Ectocarpus* filaments inside PDMS channels were similar to those previously observed in unconstrained conditions. These PDMS lab-on-chips are therefore relevant and suitable experimental devices to further study of apical growth, cell differentiation, and branching in a chemically controlled environment in combination with highly resolutive microscopy techniques.

## Material & Methods

### Chip design

PDMS was shown to be not toxic for brown algae (*Ectocarpus* sp.) (Evariste et al., 2012), where blended filaments inoculated on surfaces of different PDMS compositions grew as expected. Fig 2A shows a schematic of a PDMS device in side view. The key elements of the structure consist of the inlet, the main chamber, as well as the microchannels. The inlet has a diameter of 1.2 mm and enables filling of the chamber and loading of the samples. The height of the chamber is 100 μm to ensure sufficient availability of nutrients. Depending on the planned experiment, a design with a circular (Fig 2B & C) or triangular chamber (Fig 2D & E) can be chosen. The chamber is connected to numerous microchannels with a height of 20 μm. The reduction in height from the chamber to the channel constricts the specimen’s growth direction and, by that, allows for long-term imaging within a stable focal plane. While circular chambers connect to 44 channels with a width of 25 μm and allow for multiplex observation of the specimen, triangular chambers connected to 14 channels with a width of 40 μm to enable the detailed investigation of specific growth trajectory, e.g. undulations. The channels are separated by PDMS walls with a width of 20 μm, which is necessary to ensure proper sealing and adhesion between the PDMS device and the glass slide. Fig 2D shows a schematic of a PDMS device with a triangular chamber. At the entrance of the channels a small “funnel” guides filaments from the chamber into the channel, thereby enhancing the percentage of filaments growing into the channels. Once an entrance or channel is filled, a constriction at the end of the funnel should prevent an additional filament from entering into the already occupied channel. Additionally, the constriction reduces the risk of specimens getting flushed through the channels during the initial filling of the device. As this is especially important for the wide channels that allow undulation monitoring, only the devices with the triangular chambers were equipped with such constrictions.

**Figure 2:**
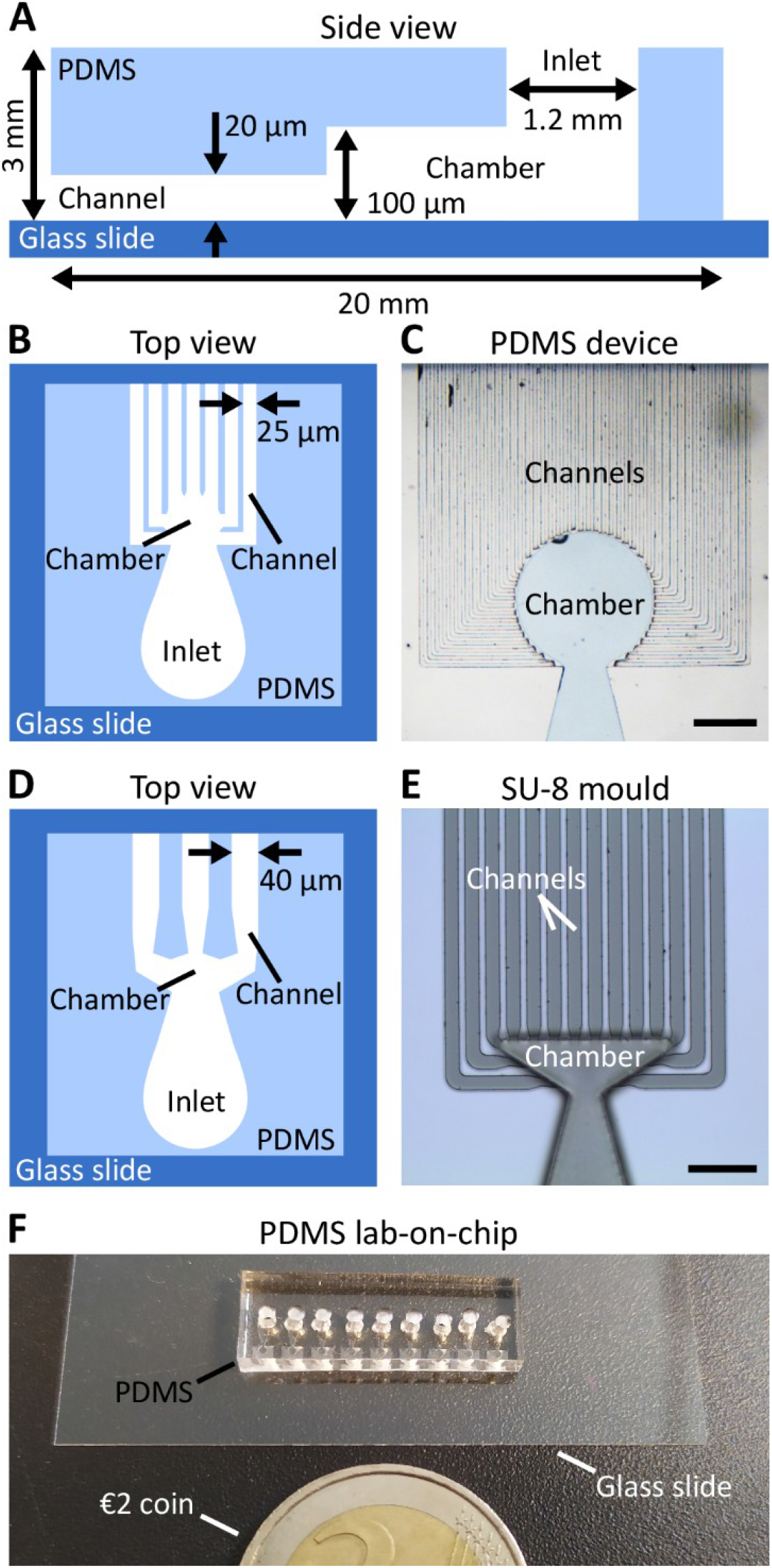
Design of the PDMS lab-on-chips. A. Schematics showing the main components (side view), including the inlet into which the algal material was injected, the chamber into which it settled, and the channels into which growth was monitored. The height of each compartment is indicated. The overall PDMS structure was fixed on a glass cover slip (0.17 μm), easing observation with an inverted microscope. B, C: Lab-on-chip made of 25 μm wide channels (top view), D, E: Lab-on-chip made of 40 μm wide channels. B, D: Schematics; C, E: Photos of the chip design (scale bars: 400 μm in C, 200 μm in E). F, G: Photos of ready-to-use lab-on-chip. F: Immersed in a Petri dish filled with seawater, G: Drying out while laying on a tip box.

The width chosen for the channels was a compromise of several criteria. On the one hand, narrower channels would not only lead to spatial constraints for *Ectocarpus* – growth cell response to constrictions is an interesting topic but was not the aim of this study – but would also complicate the fabrication process due to the reduced adhesion between the mould and the substrate. Furthermore, narrower channels would require a higher loading pressure, which could damage the samples. On the other hand, the use of wider channels bares the risk of samples passing through the microchannels rather than being collected in the chamber during loading. Overall, considering both biological features and the technical manufacturing constraints, we considered 25μm to be a compromise that would allow sufficient nutrients to be available to the algae while ensuring a mechanically robust lab-on-chip device.

As to the chamber, its geometry was adjusted to allow a sufficiently large number of connected channels for each design, while simultaneously limiting the volume available inside the chamber. Thus, it optimises the chance of obtaining filaments growing parallel to each other, a crucial factor for the efficient study of slow-growing organisms like *Ectocarpus*. However, although no direct influence on the specimen’s growth was observed in relation to changing the inlet geometry, future studies could examine the impact of additional design parameters, such as inlet height or diameter, on the specimen’s health and viability.

### Fabrication

The lab-on-chips were fabricated by double-layer photolithography in clean room environments and by mold replication techniques. Silicon wafers were cleaned with acetone, isopropanol, and deionized water prior to mold fabrication. For the layer containing the microchannels, SU-8 3025 (*Kayaku Advanced Materials*) was spin-coated on the wafers using a three-step process with a maximum angular velocity of 4500 r.p.m. to obtain a uniform photoresist layer of 20 μm. The corresponding design of the channel layer was transferred onto the resist using a mask aligner (MA6/BA6, *Süss MicroTec*). Finally, the wafers were developed (mr-Dev 600, *Micro Resist Technology GmbH*) and hard baked at 150°C for 5 minutes to improve the stability of the channels and their adhesion to the substrate. The second layer of the lab-on-chip, *i.e*. the elevated section containing the inlets as well as the main chamber, was fabricated through an additional photolithography process using SU-8 100 (*Kayaku Advanced Materials*) and a maximum angular velocity of 3000 r.p.m. to ensure a uniform 100 μm thick layer of photoresist. Fig 2E shows an optical microscopy image of a fabricated mold with a triangular chamber to be used in PDMS replication. The observed variation in the focal plane is based on the height difference between the channel layer and the (triangular) chamber layer.

Prior to PDMS casting, the fabricated structures were vapor covered with (tridecafluoro-1,1,2,2-tetrahydrooctyl) trichlorosilane (CAS 78560-45-9, *ABCR*) and heated to 120° for 10 minutes for reusability of the mold. PDMS (Sylgard 184, *Dow Corning*) with a weight ratio of 1:10 (curing agent to pre-polymer) were mixed sufficiently for 5 minutes, poured over the SU-8 mold, and degassed for 30 minutes in a vacuum chamber. To transfer the 3D pattern into the polymer, the PDMS was cured for 1 hour in an oven at 80°C. Finally, the PDMS was cut and peeled of the mold and the inlets were punched using a biopsy punch with a diameter of 1.5 mm (BP15, *Vetlab*). To chemically bond the PDMS and the cover glass, their surfaces were exposed to oxygen plasma (Femto Plasma Asher, *Diener Electronics*) for 30 seconds before brought in contact. Slight pressure was manually applied to ensure proper contact between the entirety of the lab-on-chip and the cover glass to prevent future algal specimens from growing beyond the corresponding microchannels.

For size comparison, Fig 2F presents a ready-to-use lab-on-chip with 25 μm wide channels next to €2 coin.

### Sterilisation of the chips

The chips were sterilised by 30min UV irradiation in a sterile laminar flow hood. They were then used either dried or pre-filled with sterile seawater (see below).

### *Ectocarpus* cultivation

*Ectocarpus* strain CCAP 1310/4 (also named Ec32) from the Culture Collection of Algae and Protozoa is grown in natural seawater (~ 550 mosmoles, 450 mM NaCl) supplemented by vitamins and microelements as described in Le Bail & Charrier (Le Bail and Charrier, 2013). Sporophyte filaments were produced from the germination of swimming mitospores (Charrier et al., 2008). Cultivation took place at 13°C under 12:12 Light:Dark conditions (light intensity ~ 29 μE m^−2^ s^−1^), usually on the main types of plastic and glassware. Seawater was renewed every two weeks when grown in open space.

### Chip inoculation

#### Optimisation of the loading procedure

The 1.2 mm diameter inlet was filled with seawater by applying constant pressure via a pipette mounted with a pipette tip cut to seal the inlet entrance.

Seawater is 1.3% more viscous and has a higher surface tension than pure water, making it more difficult to fill the channels. Consequently, if the applied pressure is too low, the seawater introduced into the inlet only fills the chamber but not the channels (Suppl Fig 1A and B). A solution of seawater stained with bromophenol blue was used to optimise the channel filling procedure with a pipetman. Once the protocol was established, algal material was introduced into the chamber.

#### Channel filling

The algal material was introduced into the chamber by applying a steady pressure high enough to bring the algal material just at the entrance of the channels, yet low enough to prevent the algae from being flushed out. After inoculation, the material was left still for two hours to allow the spores to attach to the chamber surface. If sporophytes were used instead of spores, they were removed after two hours. The channels were then filled with seawater by applying an adequate pipetting pressure, which was previously set by using blue seawater (Suppl Fig 1C). To confirm complete filling of the channels with clear seawater, the exit of air bubbles was monitored under the microscope. Channels in which air remained trapped (as shown in Suppl Fig 1D) were flushed several times.

Once inoculated, the lab-on-chip was transferred to a Petri dish filled with seawater so that it was fully immersed. It was cultured under standard conditions as described above, thereby allowing comparison with open space cultivation. Seawater was renewed every two days. No salt crystals were observed inside the channels during at least 3 weeks.

This procedure was repeated on 7 independent experiments summing 12 slides containing a total of 108 lab-on-chip devices. Altogether, more than 500 filaments were observed.

### Image acquisition

The Leica DMI-8 inverted microscope was used to observe calcofluor (Ex/Em : 380/475 nm) and FITC (Ex/Em : 495/520) labelling with the corresponding filters. Confocal microscopy (TCS SP5 AOBS inverted confocal microscope, *Leica*) was used to increase the z spatial resolution, compared to image acquisition by epifluorescence microscopy.

### Labelling of cellular components

Cellulose was stained by incubating the lab-on-chip in 0.01 % calcofluor white solution (fluorescent brightener 28, F-3543, *Sigma-Aldrich*) for 30 min at room temperature (RT) in the dark. The solution was injected into the channels by high pressure pipetting. Observation was performed under UV light after flushing the channels with fresh seawater at least twice and after incubation for at least 15 min at RT between each rinsing step.

Immunolocalisation of cell wall polysaccharides in the lab-on-chip was developed by adapting the protocol from Rabillé and colleagues (Rabillé et al., 2019b). The main modification consisted in fixing the algal material in 4% paraformaldehyde prepared in H_2_0 instead of seawater. All the solutions were rinsed by high pressure pipetting into the inlet, which occasionally resulted in the loss of some filaments.

## Results

### Inoculation of the chip : optimisation of the loading procedure

To study the growth of filaments from the germination stage, the lab-on-chip must be optimally inoculated with spores, as spores are the initial cells from which filaments develop. Notably, equipped with two propelling flagella, these spores are mobile and swim in a rotational movement. Therefore, the challenge consisted in having spores settle and germinate at the entrance of the channels to optimise monitoring time, rather than having them settle inside the channels, which would reduce monitoring time. We tested several parameters on two types of PDMS lab-on-chips made of parallel channels: those with width of 25 μm and those with width of 40 μm (Fig 2). The first parameter was the choice of the biological material, i.e. either swimming spores or fertile reproductive filaments. The second parameter was the location of the loaded material, i.e. either inside a wet chamber preceding dry channels (*Case 1*), or inside a wet chamber with wet channels (*Case 2*). Case 1 prevents spores from entering the channels.

#### Case 1: Loading fertile sporophytes or released spores in a dry lab-on-chip

Only the chamber and the inlet were wet and therefore the released swimming spores remained in the chamber and did not enter the channels (Movie 1). The spores bounced against the liquid/air interface. After a few hours, they attached themselves to the solid surface (glass bottom) from where they started to germinate.

#### Case 2: Loading of fertile sporophytes or spores into lab-on-chips chambers with channels filled with seawater

Spores may swim from the chamber where they were loaded into the channels. Notably, broken filaments or small pieces of filaments got stucked at the entrance of the channels (Suppl Fig 2A). By applying a higher loading pressure, spores or fertile sporophytes could even enter the channels.

Finally, an alternative combination consisted in loading non-fertile sporophytes into the chamber with channels filled with seawater. This method resulted in branches growing in the channels (Suppl Fig 2B). Because *Ectocarpus* development is reiterative, branch growth followed the same developmental process as the primary filament. Only spore germination and initial asymmetrical divisions were skipped. Therefore, this was considered as an option to have many channels filled with branches at a similar stage of development.

### Viability and developmental steps of Ectocarpus filaments within the lab-on-chip

The early development of the *Ectocarpus* sporophyte has been described in detail (Le Bail et al., 2008, 2010, 2011; Nehr et al., 2011; Rabillé et al., 2019a) and is summarised in Fig 3. When grown in open environment, the *Ectocarpus* sporophyte is composed of microscopic, branched uniseriate filaments (Fig 3A) that form a visible tuft approximatively 4 weeks after the very initial stages (Fig 3B). Sporophytic filaments growth is initiated by an asymmetrical cell division of the zygote or mitospores (Fig 3C), forming the first apical cell that continues to grow indefinitely by tip growth (Fig 3D). A few hours later, the initial cell germinates again, this time on the opposite side. It gives rise to a second apical cell which grows along the same axis as the first one but in the opposite direction. Each first apical cell gradually differentiates into spherical cells, a process requiring several days. These two processes – apical growth and cell rounding – generate a uniserial filament composed of two main cell types: elongated cells ~ 7 μm wide located at both ends of the filament and round/spherical cells ~ 15 μm wide located in the centre of the filament (Fig 3E). The sub-apical cells branch after ~ 10 days (Fig 3F) and these branches continue to grow like the primary filament. This repeated developmental program results in the overall morphology shown in Fig 3B.

**Figure 3:**
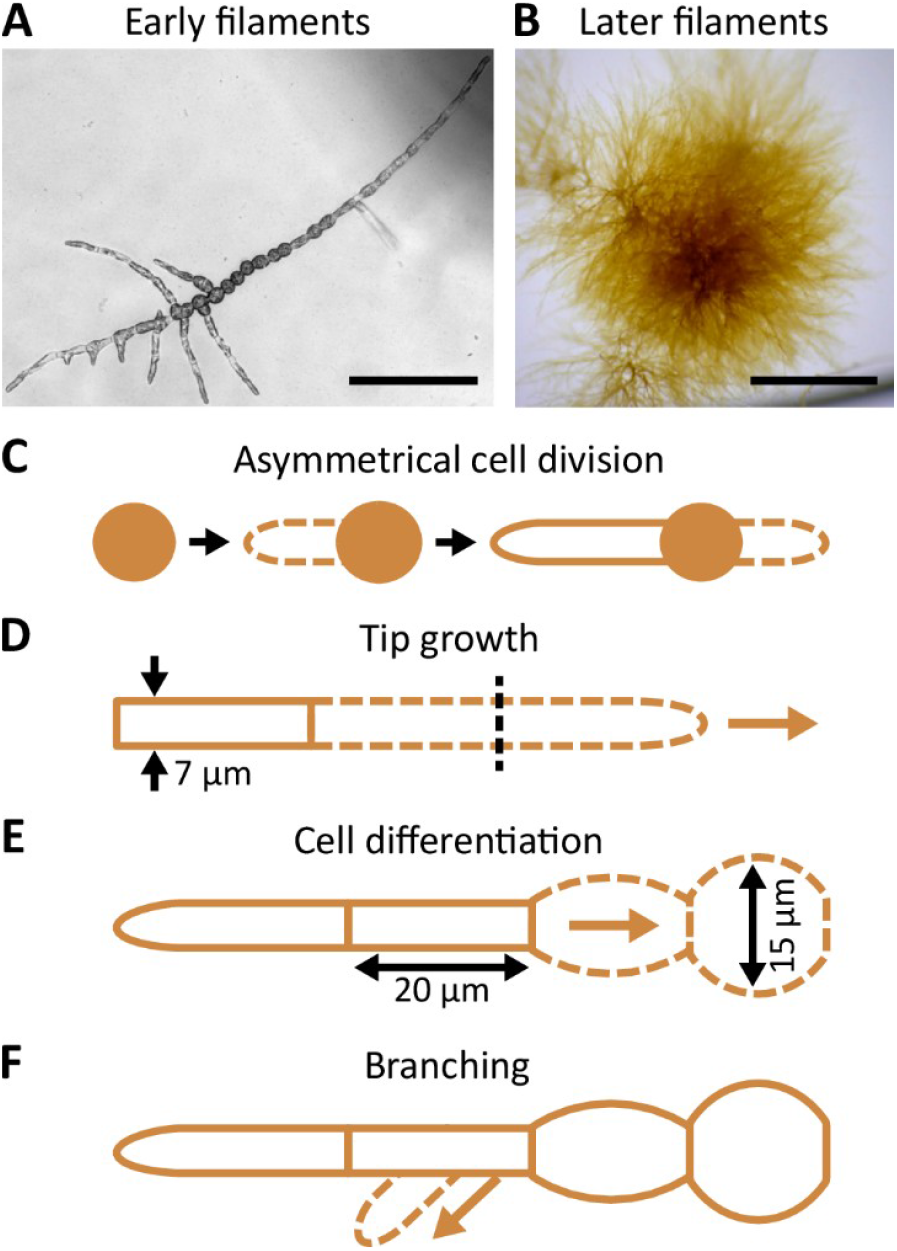
Morphology and development of *Ectocarpus* prostrate filaments. General morphology of early (A) and later (B) prostrate filaments. C-F: Schematic representation of filament growth. Zygote, mitospore or parthenogenetic gamete germinate, divide asymmetrically and produce the first, highly polarised, apical cell (C). Growth occurs at the tip of the apical cell and is indeterminate (D). The elongated cylindrical apical cells (up to 60 μm long) progressively differentiate into rounder cells (E). In the meantime, branching takes place on the shank of a sub apical cells (F). The branch re-iterates the same developmental pattern as the primary filament. Scale bars: 100 μm in A, 1 mm in B.

We investigated whether these developmental steps were preserved in the constrained lab-on-chip environment. We first monitored growth in the lab-on-chip conformation corresponding to *case 1*, i.e. wet chamber and dry channels, with the goal of constraining the spores to germinate in the chamber. We observed that *Ectocarpus* spores germinated 5 days after inoculation (Fig 4A). The rate of cell division asymmetry, i.e. the ratio of cell divisions producing unequal sized daughter cells, as depicted in Fig 3C was consistent with what has been reported in open space germination, where ~ 80 % of spores divide asymmetrically and ~ 20 % symmetrically (Le Bail et al., 2011). When grown freely in water deprived of microelements, we have previously observed that filaments tend to develop a cell sheet, meaning that *Ectocarpus* filaments shifted their development from uniaxial to bidirectional growth. This major change in body plan organisation is a sign that tip growth mechanisms have been severely impaired. In both 40 μm and 25 μm wide lab-on-chips, we did not observe such pattern, as the filaments maintained uniaxial growth throughout (Fig 4B). Calcofluor staining further showed that growth took place in the dome of the apical cell (Fig 6I), as observed in open space environments (Le Bail et al., 2008). We measured the growth rate of 22 filaments growing in 25 μm wide channels for one week and compared it to filaments growing in the open space of the same Petri dish (Fig 5A,B). Fig 5C,D showed that the overall growth dynamics was similar between the confined and free filaments, as filaments growing in the channels had an average growth rate of 2.8 μm.h^−1^ compared to 2.41 μm.h^−1^ for filaments growing in the open space (Suppl Table 1; T-test P value 0.39). In both cases, the growth rate appears to decrease with time (e.g. from 3.13 to 2.60 μm.h^−1^ in channel environment) but this it is not statistically supported (T. Test P value = 0.15). Overall, growth occurs with the same dynamics as previously reported (Rabillé et al., 2019a; 2.5 μm.h^−1^).

**Figure 4:**
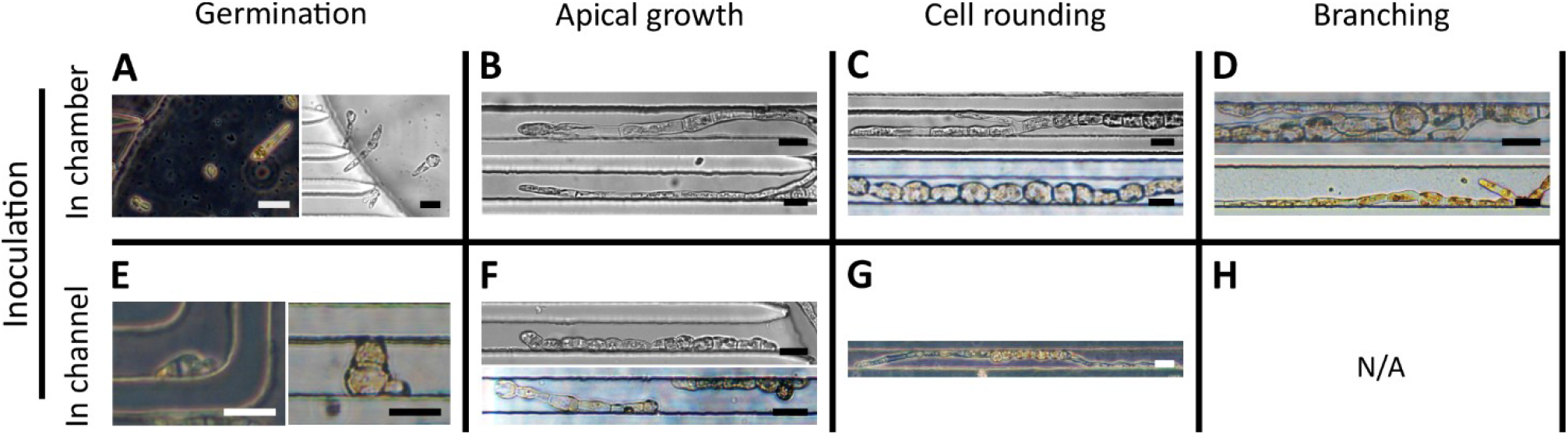
Development of *Ectocarpus* filaments in PDMS lab-on-Chips. A-D: Inoculation in the lab-on-Chip chamber only. E-H: Inoculation inside the channels. A: Germination and first division of spores. Spores divided mainly asymmetrically. B. Apical growth. C: Cell rounding. D: Branching. The lower picture is in 40 μm wide channels. E: Spore germination was impaired: it was slow and perpendicular to the channel orientation. F: Apical growth was impaired, with shorter and round apical cell and z slower growth rate. G: Cell rounding occurred in some filaments growing by tip growth. H: No branching was observed in filaments germinating inside the lab-on-chip channels. Scale bars: 25μm. Photos taken in 25 μm wide channels unless otherwise indicated.

**Figure 5:**
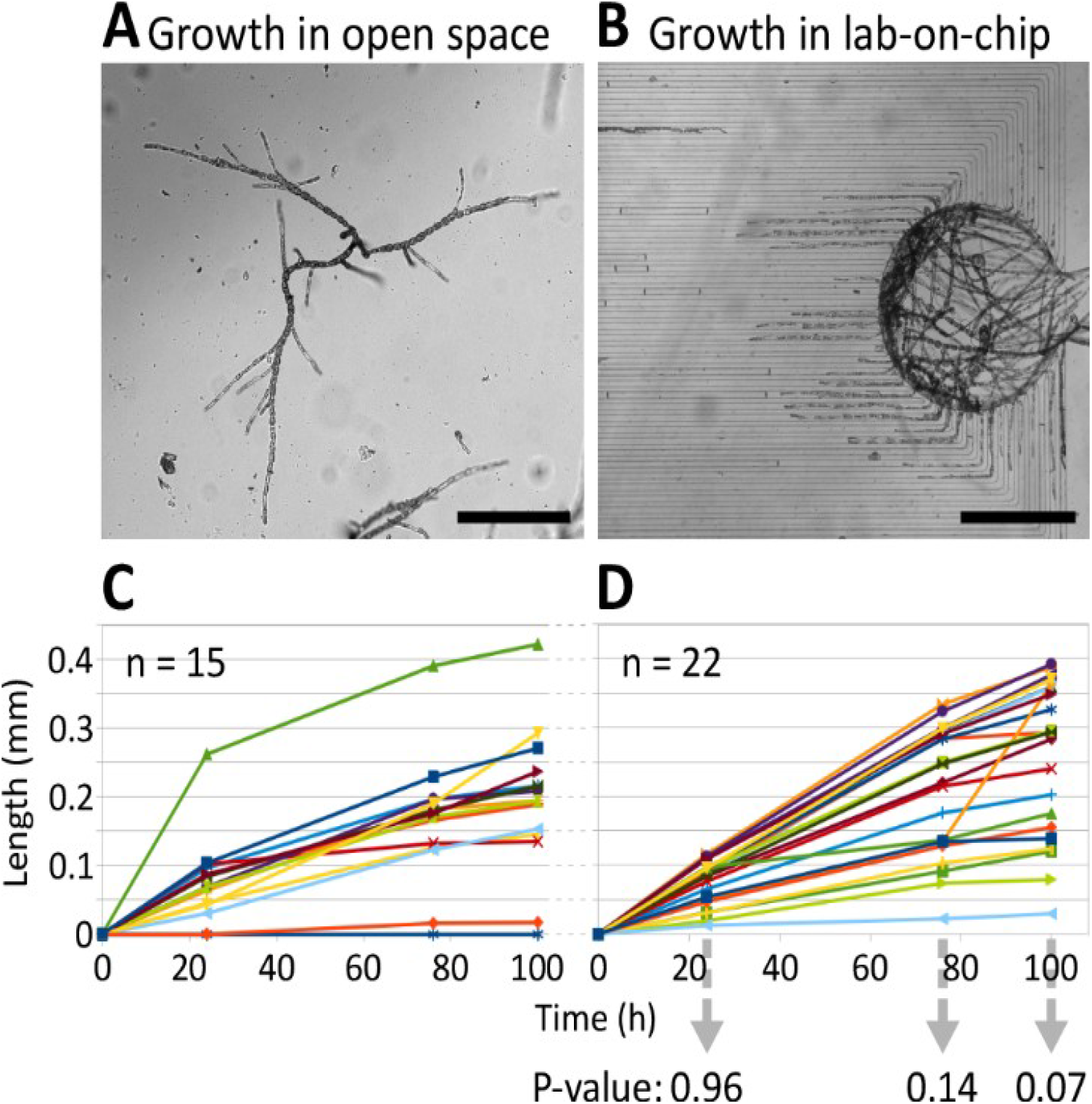
Growth rate of *Ectocarpus* filaments in lab-on-chip channels and in the open space. A, B: Photos of the growing filaments in the open space, i.e. at the bottom of the Petri dish, and inside the lab-on-chip. Scale bars = 500 μm. C, D: Growth curves over three time points: 24, 76 and 100 hours after the first recording of the initial tip position. The initial tip position was set to 0 by default so that subsequent tip positions are indicated as relative positions. P-value of T-test (two-tailed, unequal variance) is shown for each time point in the bottom of D. Sample size is indicated for each growth curve. Colours represents different filaments. Measurement data are shown in Supplementary Table 1.

**Figure 6:**
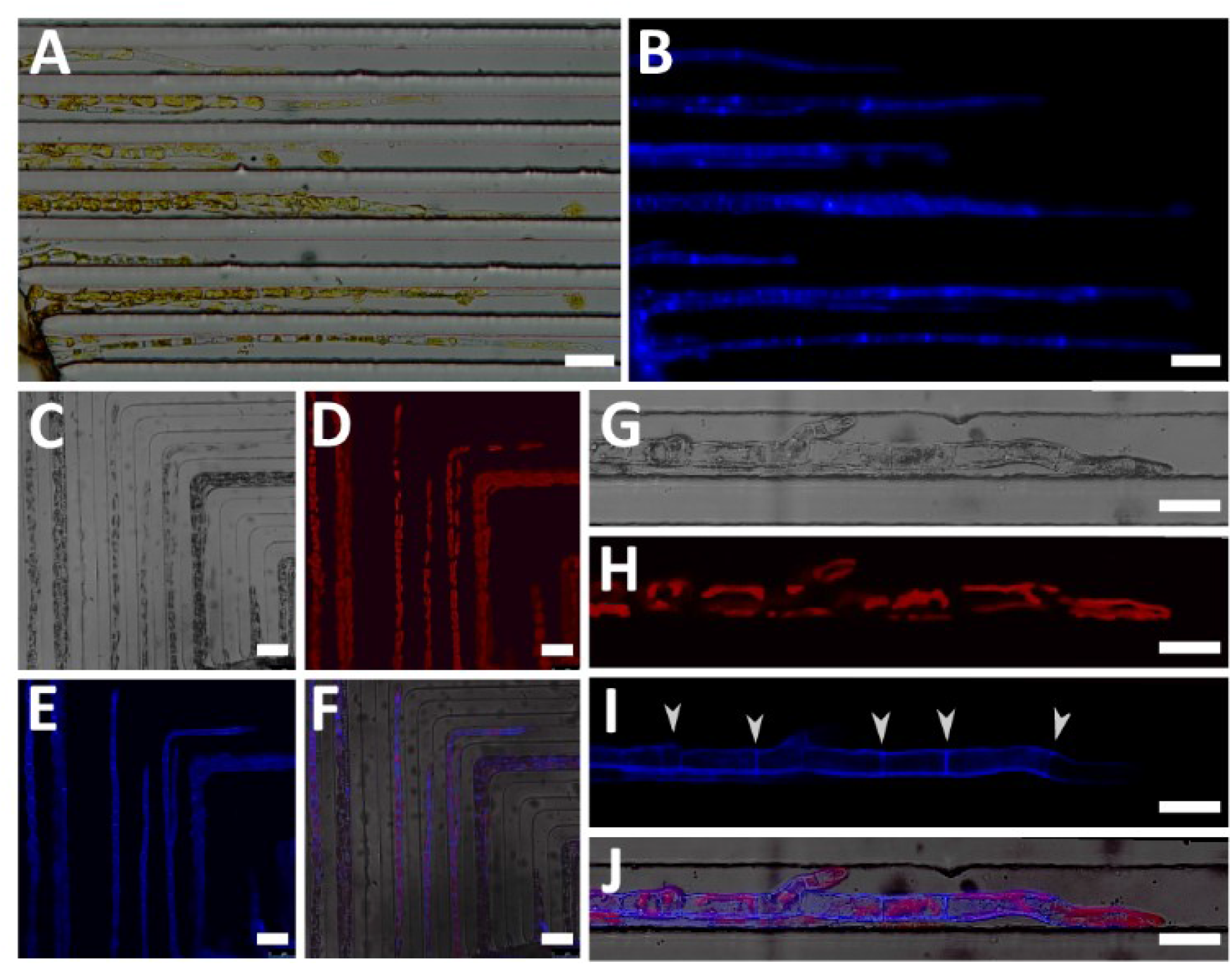
Labelling of cellulose in the cell wall of filaments growing in 25 μm wide channels. A, B: Observed in epifluorescence microscopy. A: Bright field, B: Under UV, showing cellulose present in the external cell wall in blue. A stronger signal was seen in the transversal walls. C-J: Observed with confocal microscopy. C-F: General view. C: Bright field, D: Red chloroplast autofluorescence, E: Blue calcofluor fluorescence, F: Three channels merged in. G-H: Close-up of filaments labelled with calcofluor and grown one day after washing calcofluor out. G: Bright field, H: Red chloroplast fluorescence, I: Blue calcofluor fluorescence. Transverse cell walls (white arrowheads) are clearly distinguishable. Tip is not fluorescent, and as such, displays the new grown cell wall. J: Three channels merged in. Scale bars: 50 μm in A, B; 50 μm in C-F; 25 μm in G-J.

The rounding of the cells along the filament is a function of the maturation stage of the cells and of their position along the filament. It is therefore an excellent parameter reflecting the fitness of *Ectocarpus*. Rounding of elongated sub-apical cells (Fig 3E) was observed as expected, so that spherical cells were located only in the most proximal parts of the filaments (Fig 4C). Branching, which is an additional cellular event reflecting the canonical developmental pattern described in Fig 3, occurred in filaments developing within the channels (Fig 4D, also shown in Fig 4C). Finally, we did not notice any colour change from brown to green in the chloroplasts of cells growing inside the microfluidic chip, a process observed when filaments die. Furthermore, under UV, the fluorescence intensity was similar to that of filaments growing in the open space (see Fig 6D, H below).

Therefore, when spores germinated in the chamber, the growth pattern was not affected by either the presence of PDMS, or the limitation of space that could have both impaired the quality of the seawater inside the channels and increased the mechanical stress.

Then we observed the development of spores as they germinated inside the channels (*case 2*). Spore germination was delayed in most cases, and germinated spores seemed to divide more often symmetrically (qualitative observation). In addition, the growth axis was observed perpendicular to the channel axis in some cases (Fig 4E). Apical cells were less elongated and slow growing, resulting in generally shorter filaments (Fig 4F). Cell rounding could still be observed in filaments with normal apical growth (Fig 4G). Branching was never observed in filaments whose initial spores germinated inside the channels, most likely because their developmental stage was delayed.

Similar data were obtained in PDMS lab-on-chip with 40 μm wide channels.

Altogether, when germination took place in the chamber, the developmental pattern of the filaments growing inside the channels, i.e. apical cell polarisation, growth rate and cell shape changes, was consistent with the pattern observed in filaments growing in an open space. In contrast, when the spores germinated in the channels, the developmental pattern of the growing filaments was severely impaired. Therefore, the best conformations for lab-on-chip inoculation were those that prevented spores from germinating inside the channels. Having growth monitored only from the entrance of the channels also optimised the duration of the observations along the entire length of the channel.

### Display of cellular components within the lab-on-Chip

Despite the recent development of editing approaches in *Ectocarpus* (Badis et al., 2021), expression of fluorescent reporter genes is not yet feasible. Therefore, the study of *Ectocarpus* cell biology requires us to use classical cytological approaches. Vital staining and immunolocalisation are two non-invasive techniques allowing the labelling of specific components in or at the surface of the algal cells. They are the only ones available to date because genetic transformation with visible reporter genes is not possible in this species so far. Using fluorescent calcofluor white, we first tested the staining of cellulose, a rigid polymer of β(1-4) glucose present in *Ectocarpus* cell wall at a level of ~ 10% (Charrier et al., 2019). Observed under UV epifluorescence microscopy, calcofluor uniformly labelled filaments grown in the 25 μm wide channels on both the outer and transverse cell wall (Fig 6A,B). This is in agreement with previous studies with filaments grown in open environment (Le Bail et al., 2008; Simeon et al., 2020). Similar results were obtained in 40 μm wide channels. Confocal microscopy observations provided more focused images, as shown in Fig 6C-F, which confirmed the overall and homogeneous labelling of all filaments present in the channels. Both the shape of the chloroplasts displayed by autofluorescence (Fig 6G,H) and the apical growth displayed by the new dark area at the filament tip (Fig 6I,J), confirmed that filaments were thriving in this confined environment, as previously indicated by the measurement of the growth rate.

In a second step, we labelled alginates, a polysaccharide consisting of a mixture of guluronic and mannuronic acids linked by β(1-4) bond and present in brown algal cell wall at a ratio of up to 40% (reviewed in Charrier et al., 2019). The monoclonal BAM6 that recognises mannuronan-rich alginates was used with a secondary antibody coupled to the green fluorochrome fluorescein isothiocyanate (FITC). The immunolocalisation protocol was applied in the lab-on-chip, aiming to label several filaments growing in parallel in a single channel each. Simultaneously, free organisms present in the same Petri dish were labelled and used as positive controls. In contrast to the negative control (Fig 7A-C) without primary antibody, the cell wall of both apical (Fig 7D, E) and rounding cells (Fig 7F) of labelled free-growing organisms displayed a specific signal similar to that reported in Rabillé et al. (2019b). In the 25 μm channels, *Ectocarpus* filaments also showed a strong signal in the dome of apical cells (Fig 7G, H, I, J) and on the flanks of rounding cells (Fig 7K). This pattern was as strong and specific as in the internal positive controls. The remaining cell walls of an empty plurilocular sporangium incidentally present within one of the channels also displayed significant labelling (Fig 7L). The red autofluorescence signal emitted by the chloroplasts further vouches for the healthiness of the filaments before the formaldehyde fixation step and for the cell structure-preserving properties of the immunolocalisation protocol (Fig 7A-J).

**Figure 7:**
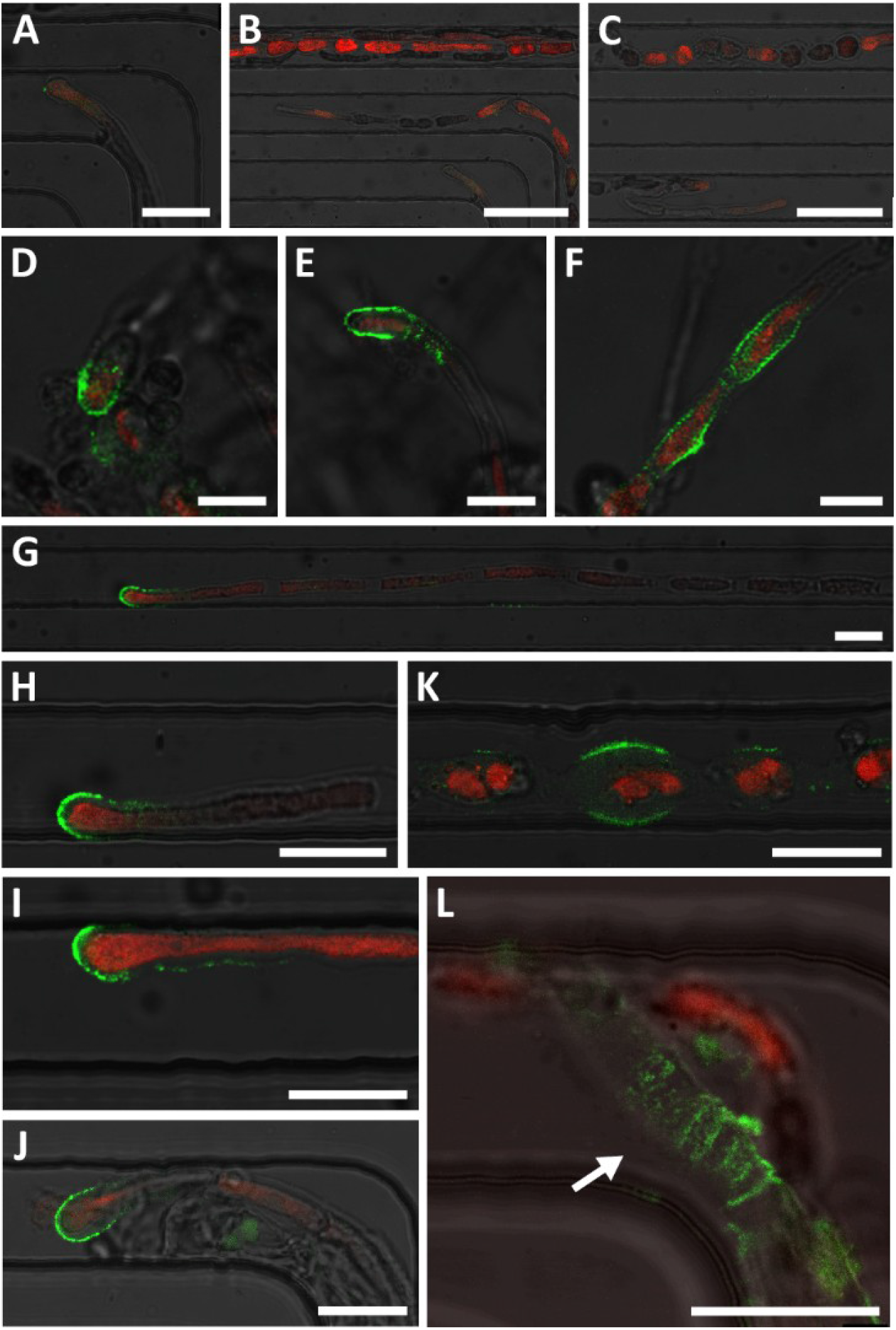
Immunolocalisation of alginate polysaccharides in *Ectocarpus* filaments, using monoclonal antibody BAM6 and observed in confocal microscopy. A, B, C: Negative controls without primary anti-alginate antibody. Both apical part and central part of several filaments are shown, devoid of green fluorescence. Only the red autofluorescence emitted by the chloroplasts was detected. D, E, F: Results on freely growing filaments in the same Petri dish. With the same image capture parameters (e.g. laser power and PMT), BAM6 labelling was observed in the dome of apical cells and in the flanks of rounded cells. G-L: Results on filaments grown in 25 μm wide channels. G-J: Dome of apical cells. K: Roundcells. L: Empty plurilocular sporangium (white arrow). Autofluorescence of chloroplasts of BAM6 labelled filaments was observed, except in L, as the cavities of the pluorilocular sporangium no longer contained spores but only cell walls remained. Scale bars: 25 μm in A, 50 μm in B, C, 10 μm in D-F, 20 μm in G-L. The specific green BAM6 signal and the red chloroplast autofluorescence signals are superimposed with the bright field signal.

Overall, these two cell biology protocols were able to label both types of cell wall polysaccharides – cellulose and alginates – that are either homogeneously distributed along the filaments or localised in a specific position, and even when the filaments were located far away from the entrance of the channels (Fig 7G).

## Conclusion

*Ectocarpus* early development is accompanied by a significant number of cellular events while its morphogenesis remains simplistic. It leads to the formation of uniseriate filaments, which grow by elongation and division of apical cells, and the progressive rounding of cells acting as the main cell differentiation process visible in this filament. Recently, we showed that tip growth relies on a different biophysical mechanism than that reported in tip growth of most cells, including the pollen tube, where the thickness of the cell wall rather than its stiffness controls growth (Rabillé et al., 2019a). This illustrated that brown algae are promising model organisms to display alternative growth mechanisms.

To further study tip growth, we tested lab-on-chips as a microfluidic device aimed at long-term monitoring of filaments. The possibility of delivering chemicals at a specific time and simultaneously to several filaments growing in-plane over a long period of time while being parallel to each other was particularly attractive. Here, we have shown that microfluidic chips with 44 parallel channels as narrow as 25 μm are suitable for studying *Ectocarpus* tip growth and dynamic cell rounding. The inoculation procedure and cell wall labelling experiments were developed such that more than 50 % of the lab-on-chip channels were filled with healthy filaments growing in-plane over more than three weeks, which can then be studied with standard cytology protocols. Similar conclusions were drawn for the moss *Physcomitrella* protonemata, where the growth rate and cell differentiation proceeded as expected (Bascom et al., 2016) – but with some impairment (Kozgunova and Goshima, 2019). However, different results were obtained with *Neurospora crassa* hyphae growing in grid or maze chips, where velocity of the apical extension was massively reduced, and their branching pattern impaired (Held et al., 2011).

In our case, the limitation was where the mitospores were loaded. While the filaments successfully entered and grew along the microchannels if the mitospores were germinated in the main chamber, mitospores loaded inside the channels were apparently hindered in their germination. However, the study of germination and especially of the first asymmetric cell division does not require any specific device since these two major cellular events can be observed in all spatial directions. Altogether, this study validates microfluidic chips to study the development and physiology of *Ectocarpus* in a confined microenvironment. It is a first step in the subsequent characterisation of tip growing mutants (Le Bail et al., 2011) and potentially life cycle mutants in this species (Coelho et al., 2011; Macaisne et al., 2017) as has been performed in *Physcomitrella* (Bascom et al., 2016) and *Neurospora* (Held et al., 2011). It will also pave the way for the study of other brown algae, especially those that develop filaments in one or both phases of their life cycle (e.g. *Sphacelaria*, and filamentous gametophytes of many brown algae including the Laminariales, the Desmarestiales and the Sporochnales; Fritsch, 1954).

## Acknowledgement

We thank Naveen Shamsudhin for discussion about the experimental setup at the onset of this project. This work was supported by ETH Zürich and, in part, by an interdisciplinary grant from the Swiss National Science Foundation (Grant Number CR22I2 166110) to B.J.N. as well as career grant from the Swiss National Science Foundation (Grant Number P2EZP2_199843) to N.F.L.

## Movie legend

<u>Movie 1</u> : The spores were filmed swimming in the chips. After ~ 30 min, they slowed down and eventually stopped moving and settled in the bottom of the device to which they remained stuck. The spores were ~ 4 μm in diameter (Lipinska et al., 2015).

## Notes

### Competing Interest Statement

The authors have declared no competing interest.

## References

Agudelo, C. G., Nezhad, A. S., Ghanbari, M., Naghavi, M., Packirisamy, M., and Geitmann, A. (2013). TipChip: a modular, MEMS-based platform for experimentation and phenotyping of tipgrowing cells. The Plant Journal 73, 1057–1068. doi: https://doi.org/10.1111/tpj.12093.

Agudelo, C., Packirisamy, M., and Geitmann, A. (2016). Influence of Electric Fields and Conductivity on Pollen Tube Growth assessed via Electrical Lab-on-Chip. Sci Rep 6, 19812. doi:10.1038/srep19812.

Badis, Y., Scornet, D., Harada, M., Caillard, C., Godfroy, O., Raphalen, M., et al. (2021). Targeted CRISPR-Cas9-based gene knockouts in the model brown alga Ectocarpus. New Phytol. doi: 10.1111/nph.17525. Epub ahead of print.

Baldauf, S. L. (2003). The deep roots of eukaryotes. Science 300, 1703–1706. doi:10.1126/science.1085544.

Bascom, C. S., Wu, S.-Z., Nelson, K., Oakey, J., and Bezanilla, M. (2016). Long-Term Growth of Moss in Microfluidic Devices Enables Subcellular Studies in Development. Plant Physiol 172, 28–37. doi:10.1104/pp.16.00879.

Bogaert, K. A., Blommaert, L., Ljung, K., Beeckman, T., and De Clerck, O. (2019). Auxin Function in the Brown Alga Dictyota dichotoma. Plant Physiol. 179, 280–299. doi:10.1104/pp.18.01041.

Bothwell, J. H. F., Kisielewska, J., Genner, M. J., McAinsh, M. R., and Brownlee, C. (2008). Ca2+ signals coordinate zygotic polarization and cell cycle progression in the brown alga Fucus serratus. Dev. Camb. Engl. 135, 2173–2181. doi:10.1242/dev.017558.

Burri, J. T., Vogler, H., Läubli, N. F., Hu, C., Grossniklaus, U., and Nelson, B. J. (2018). Feeling the force: how pollen tubes deal with obstacles. New Phytologist 220, 187–195. doi:https://doi.org/10.1111/nph.15260.

Charrier, B., Coelho, S. M., Le Bail, A., Tonon, T., Michel, G., Potin, P., et al. (2008). Development and physiology of the brown alga Ectocarpus siliculosus: two centuries of research. New Phytol. 177, 319–332. doi:10.1111/j.1469-8137.2007.02304.x.

Charrier, B., Rabillé, H., and Billoud, B. (2019). Gazing at Cell Wall Expansion under a Golden Light. Trends Plant Sci. 24, 130–141. doi:10.1016/j.tplants.2018.10.013.

Cock, J.M., Sterck, L., Rouzé, P., Scornet, D., Allen, A.E., Amoutzias, G., et al. (2010). The Ectocarpus genome and the independent evolution of multicellularity in brown algae. Nature 465, 617–621. https://doi.org/10.1038/nature09016 PMID: 20520714

Coelho, S. M., Godfroy, O., Arun, A., Le Corguillé, G., Peters, A. F., and Cock, J. M. (2011). OUROBOROS is a master regulator of the gametophyte to sporophyte life cycle transition in the brown alga Ectocarpus. Proc. Natl. Acad. Sci. U.S.A. 108, 11518–11523. doi:10.1073/pnas.1102274108.

Evariste, E., Gachon, C. M. M., Callow, M. E., and Callow, J. A. (2012). Development and characteristics of an adhesion bioassay for ectocarpoid algae. Biofouling 28, 15–27. doi:10.1080/08927014.2011.643466.

Fritsch, F.E. (1954). The Structure And Reproduction Of The Algae, Vol II, The University Press. Cambridge University Press, Cambridge.

Godfroy, O., Uji, T., Nagasato, C., Lipinska, A. P., Scornet, D., Peters, A. F., et al. (2017). DISTAG/TBCCd1 Is Required for Basal Cell Fate Determination in Ectocarpus. Plant Cell 29, 3102–3122. doi:10.1105/tpc.17.00440.

Held, M., Edwards, C., and Nicolau, D. V. (2011). Probing the growth dynamics of Neurospora crassa with microfluidic structures. Fungal Biology 115, 493–505. doi:10.1016/j.funbio.2011.02.003.

Kozgunova, E., and Goshima, G. (2019). A versatile microfluidic device for highly inclined thin illumination microscopy in the moss Physcomitrella patens. Scientific Reports 9, 15182. doi:10.1038/s41598-019-51624-9.

Le Bail, A., Billoud, B., Kowalczyk, N., Kowalczyk, M., Gicquel, M., Le Panse, S., et al. (2010). Auxin metabolism and function in the multicellular brown alga Ectocarpus siliculosus. Plant Physiol. 153, 128–144. doi:10.1104/pp.109.149708.

Le Bail, A., Billoud, B., Le Panse, S., Chenivesse, S., and Charrier, B. (2011). ETOILE regulates developmental patterning in the filamentous brown alga Ectocarpus siliculosus. Plant Cell 23, 1666–1678. doi:10.1105/tpc.110.081919.

Le Bail, A., Billoud, B., Maisonneuve, C., Peters, A., Cock, J. M., and Charrier, B. (2008). Initial pattern of development of the brown alga Ectocarpus siliculosus (Ectocarpales, Phaeophyceae) sporophyte. Journal of Phycology 44, 1269–1281.

Le Bail, A., and Charrier, B. (2013). “Culture Methods and Mutant Generation in the Filamentous Brown Algae Ectocarpus siliculosus,” in Plant Organogenesis Methods in Molecular Biology., ed. I. De Smet (Humana Press), 323–332. Available at: http://dx.doi.org/10.1007/978-1-62703-221-6_22.

Lipinska, A. P., Cormier, A., Luthringer, R., Peters, A. F., Corre, E., Gachon, C. M. M., et al. (2015). Sexual dimorphism and the evolution of sex-biased gene expression in the brown alga ectocarpus. Mol. Biol. Evol. 32, 1581–1597. doi:10.1093/molbev/msv049.

Macaisne, N., Liu, F., Scornet, D., Peters, A. F., Lipinska, A., Perrineau, M.-M., et al. (2017). The Ectocarpus IMMEDIATE UPRIGHT gene encodes a member of a novel family of cysteine-rich proteins with an unusual distribution across the eukaryotes. Development 144, 409–418. doi:10.1242/dev.141523.

Nehr, Z., Billoud, B., Le Bail, A., and Charrier, B. (2011). Space-time decoupling in the branching process in the mutant étoile of the filamentous brown alga Ectocarpus siliculosus. Plant Signal Behav 6, 1889–1892.

Michel, G., Tonon, T., Scornet, D., Cock, J. M., and Kloareg, B. (2010). Central and storage carbon metabolism of the brown alga Ectocarpus siliculosus: insights into the origin and evolution of storage carbohydrates in Eukaryotes. New Phytol. 188, 67–81. doi:10.1111/j.1469-8137.2010.03345.x.

Peyrin, J. M., Deleglise B., Saias L., Vignes M., Gougis P., Magnifico S., Betuing S., Pietri M., Caboche J., Vanhoutte P., Viovy J. L., Brugg, B. (2011). Axon diodes for the reconstruction of oriented neuronal networks in microfluidic chambers. Lab Chip 11(21), 3663–73. doi: 10.1039/c1lc20014c.

Popper, Z. A., Michel, G., Hervé, C., Domozych, D. S., Willats, W. G. T., Tuohy, M. G., et al. (2011). Evolution and Diversity of Plant Cell Walls: From Algae to Flowering Plants. Annu. Rev. Plant Biol. 62, 567–590. doi:10.1146/annurev-arplant-042110-103809.

Rabillé, H., Billoud, B., Tesson, B., Le Panse, S., Rolland, É., and Charrier, B. (2019a). The brown algal mode of tip growth: Keeping stress under control. PLoS Biol. 17, e2005258. doi:10.1371/journal.pbio.2005258.

Rabillé, H., Torode, T. A., Tesson, B., Le Bail, A., Billoud, B., Rolland, E., et al. (2019b). Alginates along the filament of the brown alga Ectocarpus help cells cope with stress. Sci Rep 9, 12956. doi:10.1038/s41598-019-49427-z.

Shamsudhin, N., Laeubli, N., Atakan, H. B., Vogler, H., Hu, C., Haeberle, W., et al. (2016). Massively Parallelized Pollen Tube Guidance and Mechanical Measurements on a Lab-on-a-Chip Platform. PLOS ONE 11, e0168138. doi:10.1371/journal.pone.0168138.

Siddique, R., Thakor, N. (2013). Investigation of nerve injury through microfluidic devices. J R Soc Interface 11(90), 20130676. doi: 10.1098/rsif.2013.0676.

Simeon, A., Kridi, S., Kloareg, B., and Herve, C. (2020). Presence of Exogenous Sulfate Is Mandatory for Tip Growth in the Brown AlgaEctocarpus subulatus. Front. Plant Sci. 11, 1277. doi:10.3389/fpls.2020.01277.

Tong, Z., Segura-Feliu, M., Seira, O., Homs-Corbera, A., Del Río, J. A., & Samitier, J. (2015). A microfluidic neuronal platform for neuron axotomy and controlled regenerative studies. RSC Advances 5(90), 73457–73466. http://doi.org/10.1039/c5ra11522a

Zhou, X., Lu, J., Zhang, Y., Guo, J., Lin, W., Norman, J. M. V., et al. (2021). Membrane receptor-mediated mechano-transduction maintains cell integrity during pollen tube growth within the pistil. Developmental Cell 56, 1030–1042.e6. doi:10.1016/j.devcel.2021.02.030.

