## Supplementary figures 1 & 2 for "Growth and immunolocalisation of the brown alga *Ectocarpus* in a microfluidic environment"

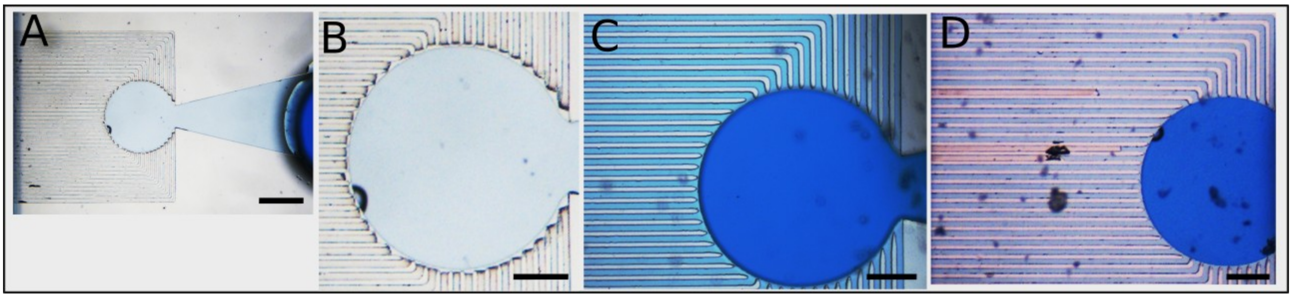


Supplementary Figure 1: Filling in the chips.
A solution of sea water with bromophenol blue was used to monitor the filling of the lab-on-chip device. A: If pipetting pressure was too low, the solution did not fill the channels and remained in the chamber (the inlet is on the right of the photo). B: Zoom-in on the channels entrance showing that the liquid did not get in. C: Fully filled channels. D: Partially filled channels. Scale bar: 500 µm in (A), 200 µm in (B-D).

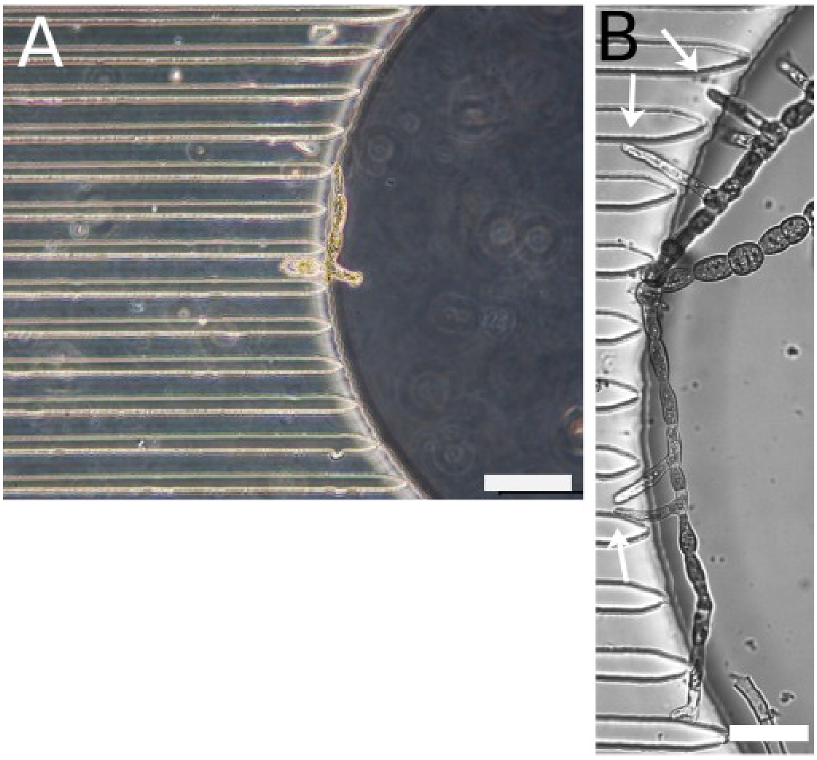


Supplementary Figure 2. Pieces of *Ectocarpus* filaments stuck in the chamber.
A: Piece of fertile sporophytes blocked at the entrance of the channels. B: Piece of a non fertile sporophyte from which branches (white arrows) emerge and grow into the channels. Scale bar: 100 µm in (A), 50 µm in (B).
